# Stability of Thin-Film Metallization in Flexible Stimulation Electrodes: Analysis and Improvement of *in vivo* Performance

**DOI:** 10.1101/644914

**Authors:** Paul Čvančara, Tim Boretius, Víctor M. López-Álvarez, Pawel Maciejasz, David Andreu, Stanisa Raspopovic, Francesco Petrini, Silvestro Micera, Giuseppe Granata, Eduardo Fernandez, Paolo M. Rossini, Ken Yoshida, Winnie Jensen, Jean-Louis Divoux, David Guiraud, Xavier Navarro, Thomas Stieglitz

## Abstract

Micro-fabricated neural interfaces based on polyimide (PI) are achieving increasing importance in translational research. The ability to produce well-defined micro-structures with properties that include chemical inertness, mechanical flexibility and low water uptake are key advantages for these devices. This paper reports the development of the transverse intrafascicular multichannel electrode (TIME) used to deliver intraneural sensory feedback to an upper-limb amputee in combination with a sensorized hand prosthesis. A first-in-human study limited to 30 days was performed. About 90 % of the stimulation contact sites of the TIMEs maintained electrical functionality and stability during the full implant period. However, optical analysis post-explantation revealed that 62.5 % of the stimulation contacts showed signs of mechanical damage at the metallization-PI interface. Such damage likely occurred due to handling during explantation and subsequent analysis, since a significant change in impedance was not observed *in vivo*. Nevertheless, whereas device integrity is mandatory for long-term functionality in chronic implantation, measures to increase the bonding strength of the metallization-PI interface deserve further investigation. We report here that silicon carbide (SiC) is an effective adhesion-promoting layer resisting heavy electrical stimulation conditions *in vivo*. Optical analysis of the new electrodes revealed that the metallization remained unaltered after delivering over 14 million pulses *in vivo* without signs of delamination at the metallization-PI interface. Reliable adhesion of thin-film metallization to substrate has been proven using SiC, improving the potential transfer of micro-fabricated neural electrodes for chronic clinical applications.

## 1. Introduction

In recent years, micro-fabricated devices initially developed for use as high resolution tools in neuroscientific animal research have entered translational research in human clinical trials. To interface the nervous system electrically, various designs and approaches have been proposed (Ordonez et al. 2012b). Devices and techniques vary in their degree of invasiveness and selectivity (Navarro et al. 2005), ranging from electrodes that lay superficially on or around the target tissue, for example electrocorticogram (ECoG) electrodes (Rubehn et al. 2009) or cuff electrodes (Naples et al. 1988), to more invasive devices that penetrate the tissue, such as intracortical microelectrode arrays (MEA) (Campbell et al. 1991; Rousche and Normann 1998). or intraneural Utah slanted electrode arrays (USEA) (Branner et al. 2004). Electrode materials and coatings need to be selected according to the manufacturing technology (Hassler et al. 2011; Navarro et al. 2005) and target application (Cogan 2008). Among the possible polymers utilized for miniaturized nerve interfaces, polyimide (PI) comes along with certain benefits (Stieglitz et al. 2000b). In comparison to silicone rubber, it is possible to achieve device thicknesses, about 10 times thinner, which help to reduce the insertion trauma for intraneural and intracortical applications. Further benefits are high chemical inertness and flexibility compared to silicon and metal wires and in case of the PI type utilized within this study, the water uptake is with 0.5 % very low (Stieglitz et al. 2000b). Moreover, fabrication using standard photolithographic processes is feasible. *In vitro* characterization indicated the possibility of using PI as substrate material for chronic application (> 30 days) (Rubehn and Stieglitz 2010). *In vivo* chronic demonstration of PI devices came when Rubehn et al. showed chronically stable recording of brain signals with a micro-fabricated 252-channel ECoG *in vivo* in Rhesus monkey (Rubehn et al. 2009). Given the possibility of chronic recording stability, the next challenge emerged: Is chronically stable stimulation possible with micro-fabricated PI electrodes?

Electrical stimulation through thin-film structures presents several challenges not seen when the devices are used for recording. The stimulation contact site must be able to transmit electrical current at a sufficient level to activate the nerve fibers without eroding or damaging the contact or electrode structure. For micro-stimulation through contacts of sizes in the 10s of µm, the safe current density limits become a challenge due to the small area of the contact (Rossini et al. 2010). For our application, i.e. multi-channel micro-stimulation of human peripheral nerves, the relatively large size of the nerve bundle further increases the challenge since penetration of the current into the larger nerve bundle requires larger currents as compared to those needed for smaller nerve bundles in small animal models. We converged to an intrafascicular approach in this particular application, since such a device places the electrode contacts within the nerve bundle, and thus achieves a more focal activation of nerve fibers while minimizing the current needed for activation. However, this approach requires multiple sites of stimulation within a given nerve fascicle or in several nerve fascicles to provide different types of sensations and cover the largest possible sensory field. Our group developed the transverse intrafascicular multichannel electrode (TIME) (Boretius et al. 2010), to increase the spatial selectivity of peripheral nerve stimulation electrodes, compared to cuff and longitudinal intrafascicular electrodes (LIFEs) (Badia et al. 2011a). The TIME is implanted perpendicularly to the axis of the nerve fascicle, in order to address as many fascicles as possible and thus to increase the stimulation selectivity. To decrease the probability of nerve damage, PI was chosen as substrate material due to its mechanical properties and manufacturing technologies, that allowed for thin, shallow and flexible devices. Before application *in vivo*, comprehensive *in vitro* studies were conducted. Stimulation with one billion pulses using constant charge of 60 nC per phase (200 µs * 300 µA) in phosphate buffered saline (PBS) solution revealed no significant change in the electrochemical properties (Boretius et al. 2012; Boretius 2013). Neither scanning electron microscopy (SEM) nor atomic force microscopy (AFM) revealed any morphological changes.

After comprehensive preclinical evaluation of the TIME in small and large animal models (Badia et al. 2011b; Badia et al. 2011a; Harreby et al. 2015; Kundu et al. 2014), a first-in-human study was conducted with an implantation period of 30 days, that has been chosen according to European medical device regulations. The aim was to stimulate the median and ulnar nerves in the stump of an arm amputee to evoke sensations at high spatial resolution, with the aim to reduce phantom limb pain and to deliver sensory feedback from a sensorized hand prosthesis governed via electromyographic voluntary commands from the stump residual muscles. The outcome of the clinical trial was very satisfying, showing the possibility of delivering near-natural sensory information to the patient by stimulation of the peripheral nerves during real-time decoding of grasping objects of different physical properties with the hand-prosthesis. It was possible for the patient to modulate the grasping forces of the hand-prosthesis and to recognize objects with different shape and compliance even when visual and auditory inputs were avoided (Raspopovic et al. 2014). Moreover, the amputee was able to discriminate with artificial fingertips the spatial coarseness (Oddo et al. 2016) of a moving surface.

Within this study, we investigated the integrity of the thin-film TIMEs implanted in the afore-mentioned first-in-human study, regarding the electrical properties and the mechanical integrity post explantation. Although the *in vivo* impedance measurements were excellent and patient responses to electrical stimulation via the TIMEs were stable up to the last day of implantation, we observed mechanical failure of the thin-film metallization, like crack formation and delamination after explantation, although none of the preceding *in vitro* and *in vivo* experiments indicated any of these kind of malfunctions. Consequently, we intended to improve the mechanical stability of the thin-film metallization, in order to ensure good performance over much longer periods of *in vivo* implantation. For this purpose, we introduced silicon carbide (SiC) as adhesion promoting layer between PI and platinum, according to our previous *in vitro* research (Ordonez et al. 2012a; Ordonez et al. 2012b; Ordonez 2013). The *in vivo* validation was performed in a small animal model. Optical analysis confirmed that the mechanical integrity of the thin-film electrodes was improved significantly using SiC as adhesion promoting layer.

## 2. Materials and Methods

The TIME implants consist of a PI-based thin-film electrode array, attached to a helically wound cable (MP35N) via an interconnecting ceramic, that terminates in a commercially available connector. The following paragraphs describe the design and the analysis of the human implant as well as the development of the improved generation that has been tested in an animal model.

### 2.1. Design of the TIME Thin-Film Layout

The version TIME-3H (details in (Boretius et al. 2012)), used in the first-in-human study sub-chronically, contained 18 channels, of which 16 correspond to stimulation contact sites (only 14 connected, due to the limited number of channels in the connector) and two ground contacts. After folding, the electrode displayed seven connected stimulation contact sites on the left and seven on the right, named L1 to L7 and R1 to R7 respectively. The same applied for the two ground contacts, L GND and R GND. The stimulation contact sites had a circular shape with a diameter of 80 µm. The large ground sites exhibited a rectangular shape with an area of 1.00 x 0.25 mm² each. Platinum tracks and pads were sandwiched between polyimide as substrate and insulation layer. At the stimulation and ground contact sites, the platinum layer was coated by iridium, subsequently covered by a sputtered iridium oxide film (SIROF) (Figure 1a top) and opened via reactive ion etching (RIE). The thin-film version TIME-3H-SiC used the same design, but with a different layer setup in order to investigate SiC as adhesion promoting layer (Figure 1a bottom).

**Figure 1.**
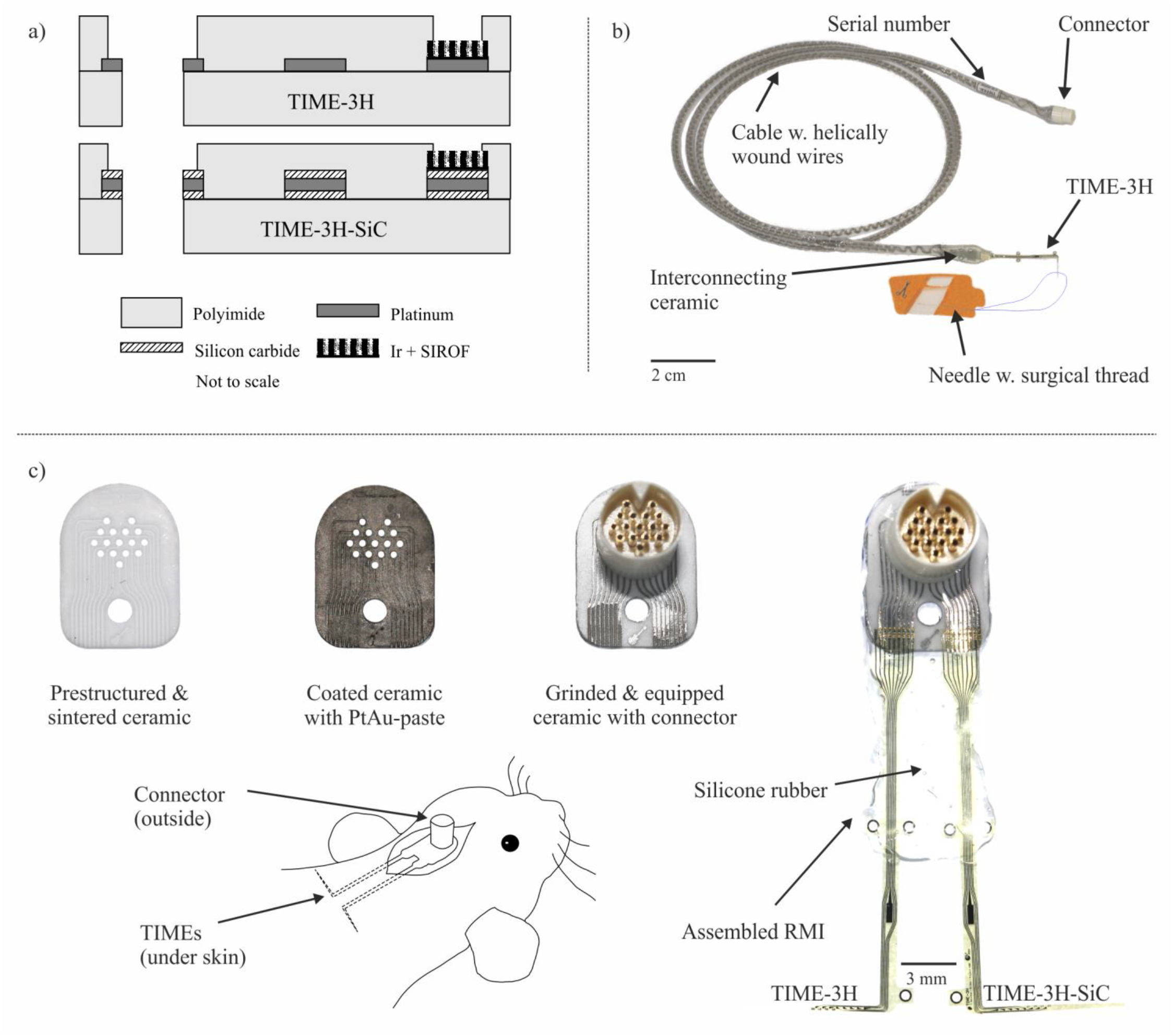
Overview of utilized implants in human and animal trials. For the implants used in the small animal model the layer setup was changed compared to the human clinical trial (a). SiC was added to the metallization to increase adhesion between PI and metallization. Assembled TIME-3H implant for sub-chronic first-in-human clinical trial (b). The implant consisted of the PI-based thin-film electrode with an incorporated surgical needle with a suture loop. The electrode was attached to the interconnecting ceramic via the MFI technique (Stieglitz et al. 2000a). A 40 cm long cable composed of 16 helically wound MP35N wires in a silicone rubber hose was soldered to the interconnecting ceramic and on the opposite end to the Omnetics connector. In order to investigate SiC as adhesion promoting layer a rodent model implant (RMI, c) was designed. The green tape ceramic was laser structured with a Nd:YAG-Laser using λ = 1064 nm wavelength. After sintering, the walls of the vias and the tracks where filled with PtAu-screen-printing paste and burned. Excessive paste was grinded and the ceramic was equipped with an Omnectics connector. Finally, half a TIME-3H and half a TIME-3H-SiC were attached utilizing the MFI technique and casted at crucial points with medical grade silicone rubber. The implant was fixed with screws and cement on the rodent skull and the TIMEs were placed underneath the skin.

### 2.2. Cleanroom Fabrication of Thin-Film Electrodes

The thin-film electrode arrays of the TIME implants were fabricated using standard processing of microelectromechanical systems (MEMS) within a clean room environment. Both thin-film electrode versions used PI (U-Varnish S, UBE Industries, LTD., Tokyo, Japan) as substrate and insulation material. The thin-film electrodes for the TIME-3H was fabricated according to previously published processes (Boretius et al. 2010; Boretius et al. 2012). Further details can be found in the supplementary materials.

For the TIME-3H-SiC – in order to evaluate SiC as adhesion promoter – a 50 nm layer of SiC was deposited using plasma-enhanced chemical vapor deposition (PECVD, PC310 reactor, STS Surface Technology Systems plc, Newport, UK) before evaporation of a 300 nm layer of platinum (Leybold Univex 500, Leybold Vacuum GmbH, Cologne, Germany) (Figure S2b). Afterwards, another adhesion layer of 50 nm SiC was deposited via PECVD on the platinum (Figure S2b1). All other fabrication process steps were the same for both TIME versions.

Following the last fabrication step, the thin-film electrodes were pulled off the silicon wafer with a pair of forceps (Figure S2f) for assembly of the implants.

### 2.3. Implant Assembly

The TIME-3H implants (hereafter called TIME) were assembled out of four sub-modules (for details see (Boretius et al. 2012) and Figure 1b). First, the thin-film part containing the stimulation contacts and the ground contacts to close the electrical circuit during stimulation. Second, a screen-printed interconnecting ceramic to mechanically and electrically connect the thin-film electrode to the third part, a 40 cm long cable. The fourth module was a commercially available connector (NCP-16-DD, Omnetics Connector Corporation, Minneapolis, USA).

The TIME implants were fabricated under an accredited quality management system (ISO 13485) at the Laboratory for Biomedical Microtechnology of the Albert-Ludwig-University of Freiburg, Freiburg, Germany. Before implantation, each stimulation and ground contact site of the TIME was hydrated in order to increase the charge injection capacity (Boretius and Stieglitz 2012), and tested on functionality. Afterwards, each TIME was washed, wrapped in sterile bags, labelled and steam sterilized at 121 °C and 2 bar for 21 minutes.

Before transferring thin-film electrodes enhanced with SiC as adhesion promoting layer into human implants, an *in vivo* verification concerning function and validation regarding stability was performed. For this, a rodent model implant (RMI) was developed, which allowed to implant and deliver stimulation with a TIME-3H and a TIME-3H-SiC simultaneously in the same animal (Figure 1c).

For this rodent model, a high temperature co-fired ceramic (HTCC) made of aluminum oxide (HTCC 44000, ESL Europe, Agment Ltd., Reading, UK) served as carrier substrate. Details of this process can be found in Fiedler et al. (Fiedler et al. 2013). After lamination of four layers of the not-sintered HTCC Al_2_O_3_ substrate, vias and tracks were laser structured with a Q-switched Nd:YAG-Laser (λ = 1064 nm, DPL*Genesis* Marker, cab Produkttechnik GmbH & Co. KG, Karlsruhe, Germany). Afterwards, the substrate was sintered according to the datasheet in a high temperature furnace for 2 h at 1500 °C. The walls of the vias and the complete tracks were filled with PtAu screen-printing paste (5837-G, ESL Europe, Agment Ltd., Reading, UK) and fired according to the datasheet in a furnace (PEO-601, ATV-Technologie GmbH, Vaterstetten, Germany). Excessive screen-printing paste was removed by grinding.

The design was chosen to fit a commercially available connector (NCS-DD-16, Omnetics Connector Corporation, Minneapolis, MN, USA). This connector was soldered from the backside of the substrate after removing the connector pins. Excessive solder was removed by grinding. The design of the tracks was customized according to the microflex interconnection technique (MFI) structure (Stieglitz et al. 2000a).

For validating both TIME-3H and TIME-3H-SiC, at the same time in the same animal, the thin-film electrodes were cut at the middle line, so it was possible to attach both thin-film versions next to each other (Figure 1c). After bonding using the MFI technique, the thin-film electrodes were stabilized mechanically with epoxy (UHU Endfest Plus 300, Bolton Adhesives, Rotterdam, Netherlands). To prevent shorts and corrosion of metal parts, the unprotected parts of the implant were casted in medical grade silicone rubber (MED-1000, NuSil Technology LLS, Carpinteria, CA, USA) (Figure 1c). In total we used four devices in four rats.

### 2.4. Analysis of TIME-3H from Sub-Chronic First-in-Human Clinical Trial

The focus of this work was the evaluation of the TIME-3H implant of a sub-chronic first-in-human clinical trial which results and outcome have been described in detail previously (Raspopovic et al. 2014). The patient was a 36-year-old male, suffering from a traumatic transradial amputation of the left arm ten years before implantation. The subject was implanted with TIME 1 and 2 in the median nerve (proximal and distal) and TIME 3 and 4 in the ulnar nerve (proximal and distal). The implantation surgery was conducted on 26^th^ of January 2013 at the Polyclinic A. Gemelli at the Catholic University, Rome, Italy after obtaining the ethical approval. The patient signed an informed consent form before the beginning of the study. For all experiments conducted in the study, the respective EU guidelines and regulations were considered.

At the end of the 30 days’ limit of the clinical trial, the implants were removed according to the clinical protocol. The patient underwent a six months’ follow-up after finishing the clinical trial and did not reveal any side-effects, neither objective nor subjective.

### 2.5. Electrical Characterization of the TIMEs *in vivo* (Human)

Data during *in vivo* experiments were acquired by the STIM’nD stimulator (max. output current 5 mA at a maximum of 20 V) and associated software provided by INRIA (Montpellier, France). The impedance of each stimulation contact site and the ground contacts of the four implanted TIMEs was estimated using an indirect method that is well established in clinical neural implants. For this purpose, a very small controlled-current stimulus (balanced biphasic rectangular pulses of I = 40 µA and t = 300 µs per phase) was generated using each active contact site separately (cathodic phase first) and one ground contact site. A resistor of R = 1 kΩ was inserted in series with the TIME implant to monitor the current. While generating the stimulus, the voltage-drop between cathode and anode as well as the voltage-drop at the resistor were measured (insulated NI6218, National Instrument, Austin, Texas, USA) with an acquisition frequency of 100 kSs^−1^. The resistance part of the impedance of each stimulation contact site versus each of the two ground contacts (left and right) as well as between the two ground contacts of each TIME was determined as the ratio of the voltage excursion at the end of the cathodic phase divided by the current amplitude of the stimulation pulse.

Despite electrodes with an impedance up to 250 kΩ can be driven by the stimulator, we included a technical safety margin in our considerations and defined |Z| = 150 kΩ as technical limit in our investigations.

The impedance measurements were performed intra-operatively to verify the correct position of the TIME, and immediately after the surgical implant (day 0). The procedure was repeated on days 2, 11, 17 and 30 in order to track impedance changes during the course of the clinical trial.

### 2.6. Animal Trials in a Rodent Model

Four RMI devices described above were built up, characterized and prepared for implantation in order to analyze the long-term stability under chronic stimulation conditions in a rodent model. All experimental procedures performed were approved by the Ethical Committee of the Universitat Autònoma de Barcelona in accordance with the European Communities Council Directive 2010/63/EU, and in compliance with the NIH Guide for Care and Use of Laboratory Animals. Surgical procedures were performed in four Sprague-Dawley rats under pentobarbital anesthesia (40mg/kg i.p.). The devices were mounted on the skull and fixed with three screws and dental cement. The thin-film part was carefully located underneath the skin of the dorsum of the rat (Figure 1c). The subcutaneous location was preferred to an intraneural implant for two main reasons: it allowed to have one TIME-3H and one TIME-3H-SiC implanted under exactly the same environment conditions, and it did not induce discomfort to the animal over hours of continuous stimulation. The wound was closed around the connector, allowing the experimenter connecting the devices to an external stimulator setup (STIM’nD, INRIA, Montpellier, France & Axonic, Vallauris, France).

The RMIs were used to deliver stimulation through both TIME-3H and TIME-3H-SiC arms contact sites. Two stimulation sessions of 90 minutes per day were performed. Rectangular, charge-balanced stimulation pulses were applied with an intensity of I = 140 µA, a pulse-width of p_w_ = 150 µs and a frequency of f = 50 Hz. Due to time and equipment limitations only 5 contacts (both GNDs and three contact sites) were stimulated on every RMI. The RMIs were maintained implanted up to 1 month. A median stimulation period of 78 h per animal (representing 14 million pulses) was performed, which corresponds to about more than two months of stimulation with the clinical protocol of planed chronic human clinical trials.

### 2.7. Statistical Analysis

The data acquired *in vivo* was longitudinal and unbalanced for each time point (days). Moreover, the data was not normal distributed and the variance was not homogeneous. Therefore, we applied a linear mixed effect model for statistical analysis using the software R (version 3.5.2, The R Foundation for Statistical Computing, Vienna, Austria) and RStudio (version 1.1.463, RStudio Inc., Boston, MA, USA), which is robust to non-normality and unbalanced data sets. A paired-sample Wilcoxon signed rank test (OriginPro 2019, Version 9.6.0.172, OriginLab Corporation, Northampton, MA, USA) with a Bonferroni correction was applied in case of significant differences.

### 2.8. Optical Analysis

The optical analysis of both, the stimulation and the ground contact sites was done after explantation using light microscopy (Leica DM400M, Leica Microsystems GmbH, Wetzlar, Germany) – partially with polarization filters, to enhance visibility of surface irregularities and delamination. The view from the bottom side of the electrodes was of particular interest, as hereby delamination was better visible.

In order to analyze the properties of the layer setup, scanning electron microscopy (SEM) in combination with focused ion beam (FIB, Zeiss Auriga 60, Carl Zeiss AG, Oberkochen, Germany) was utilized to gather cross-sections of high resolution overviews of the contact sites and the thin-film metallization.

After explantation of the TIMEs from the human clinical trial, 40 of 64 stimulation contacts (including not electrically connected contacts) and 5 out of 8 ground contacts were available for optical analysis. Concerning the RMIs, 24 stimulation contacts and 4 ground contacts were available for TIME-3H electrodes and 24 stimulation contacts and 3 ground contacts in case of the TIME-3H-SiC.

A contact was declared as mechanically damaged during the optical analysis if the platinum thin-film metallization delaminated from the PI and lifted, no matter if it vanished or not. Crack formation without delamination was not considered as damage.

## 3. Results

### 3.1. Electrical Characterization of the TIMEs *in vivo* (Human)

The impedance of each single stimulation contact site and ground contact was recorded in the sub-chronic human clinical trial for all implants. The stimulation contacts were measured versus L GND (Figure 2) and R GND (supplementary material). The same was performed for the GNDs. The impedance increased during the 30 days as expected due to the foreign body reaction (Figure 2). TIME 1 exhibited on day 0 (just after implantation) large variability, with a median impedance of 84 kΩ ± 75 kΩ, that decreased during the course of the trial. The L GND contact was stable with an impedance of 25 kΩ (Figure 2, upper left). TIME 2 had increasing impedance from 31 kΩ to 91 kΩ together with increasing variation from 9 kΩ to 67 kΩ. On the last day, six contacts had an impedance higher than 150 kΩ or even not measurable and were above our predefined threshold. The L GND contact showed stable impedance lower than 27 kΩ throughout the clinical trial (Figure 2, upper right). The impedance of the contact sites of TIME 3 remained stable throughout the study, at around 60 kΩ, except at day 0, when the impedance was 19 kΩ. The impedance of the L GND was not measurable on day 30 (Figure 2, lower left). The impedances of the active sites on TIME 4 increased from 20 kΩ ± 3 kΩ on day 0 to 89 kΩ ± 8 kΩ on day 30. The impedance of the L GND in TIME 4 increased along the trial (Figure 2, lower right) from 4 kΩ to 31 kΩ. In total, the mean impedance of all stimulation contact sites increased from 39 kΩ ± 46 kΩ to 78 kΩ ± 33 kΩ, measured against the L GND of the respective device.

**Figure 2.**
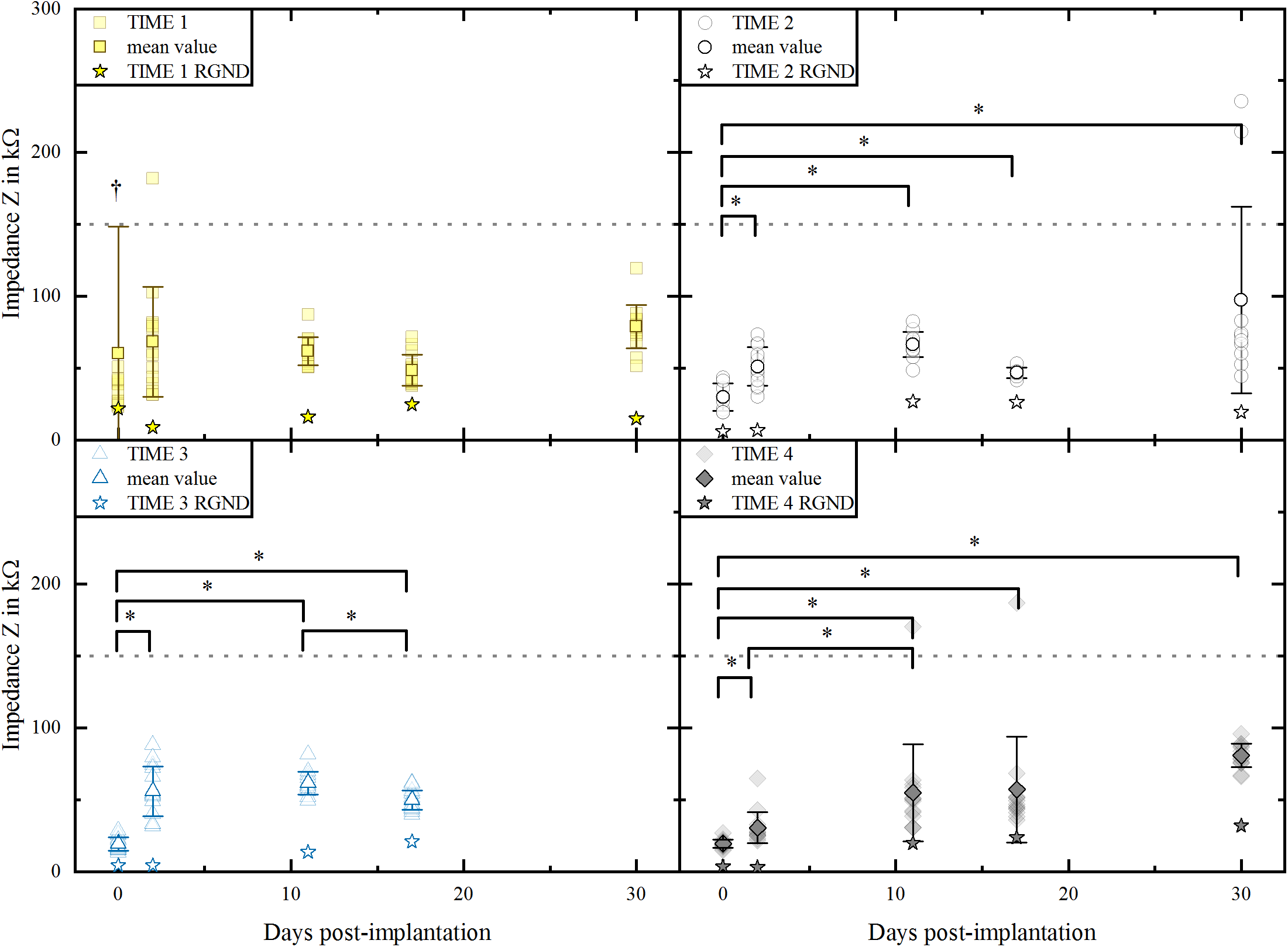
Impedance progression of four sub-chronically implanted TIME-3H implants. The impedances of the single active contact sites were measured versus the L GND of the respective implant. The measurements of each contact site are transparent, whereas the mean values are in bright color. The stars indicate the impedance of L GND, measured versus R GND. Electrical non-functionality is marked at 150 kΩ. The impedances of TIME 1 were statistically not different between each time point (linear mixed effect model; p = 0.263). There was statistically significant difference between the impedances of specific days (especially day 0 compared to the other days) of the implants TIME 2, TIME 3 and TIME 4 (paired-sample Wilcoxon signed rank test with Bonferroni correction; p < 0.01). An increase in impedance during the first month *in vivo* was expected due to foreign body reaction, as shown by other groups (Rossini et al. 2010; Williams et al. 2007; Wurth et al. 2017).

No significant difference was observed in the impedance of TIME 1 versus L GND (p > 0.05). For the other implants, there was a significant increase in the impedance over time (p < 0.05, paired-sample Wilcoxon signed rank test with a Bonferroni correction) (Figure 2). The impedance of TIME 2 and TIME 3 stabilized until the end of the clinical trial. TIME 4 was stable until day 17, but increased significantly (p < 0.01) on the last day.

Data recorded with R GND was similar to that of L GND and can be found in the supplementary material.

### 3.2. Optical Analysis of Explanted TIME Implants (Human)

Optical analysis of the explanted electrode contact sites focused on thin-film integrity revealed that 62.5 % of the available contact sites and all the available ground contacts were mechanically damaged. Delamination of the thin-film metallization always appeared between the PI and the platinum interface as adhesion failure, never between platinum and SIROF. No cohesion failure was observed.

The R GND contact of the implant TIME 3 delaminated almost entirely (Figure 3), in accordance to an electrical failure (Figure S4). The metallization lifted from the PI. The platinum showed a jagged breaking line. The contact site L6 on TIME 2 was electrically functional until the end of the clinical trial, and there was no sign of mechanical failure (Figure 3), only some dried salt crystals. A more detailed cross-section view indicated a crack formation in the SIROF, near the PI-substrate-metal transition (Figure 3, black arrow). L8 on the implant TIME 2 was not electrically connected and consequently not stimulated during implantation, however, the delamination of the thin-film metallization was severe.

**Figure 3.**
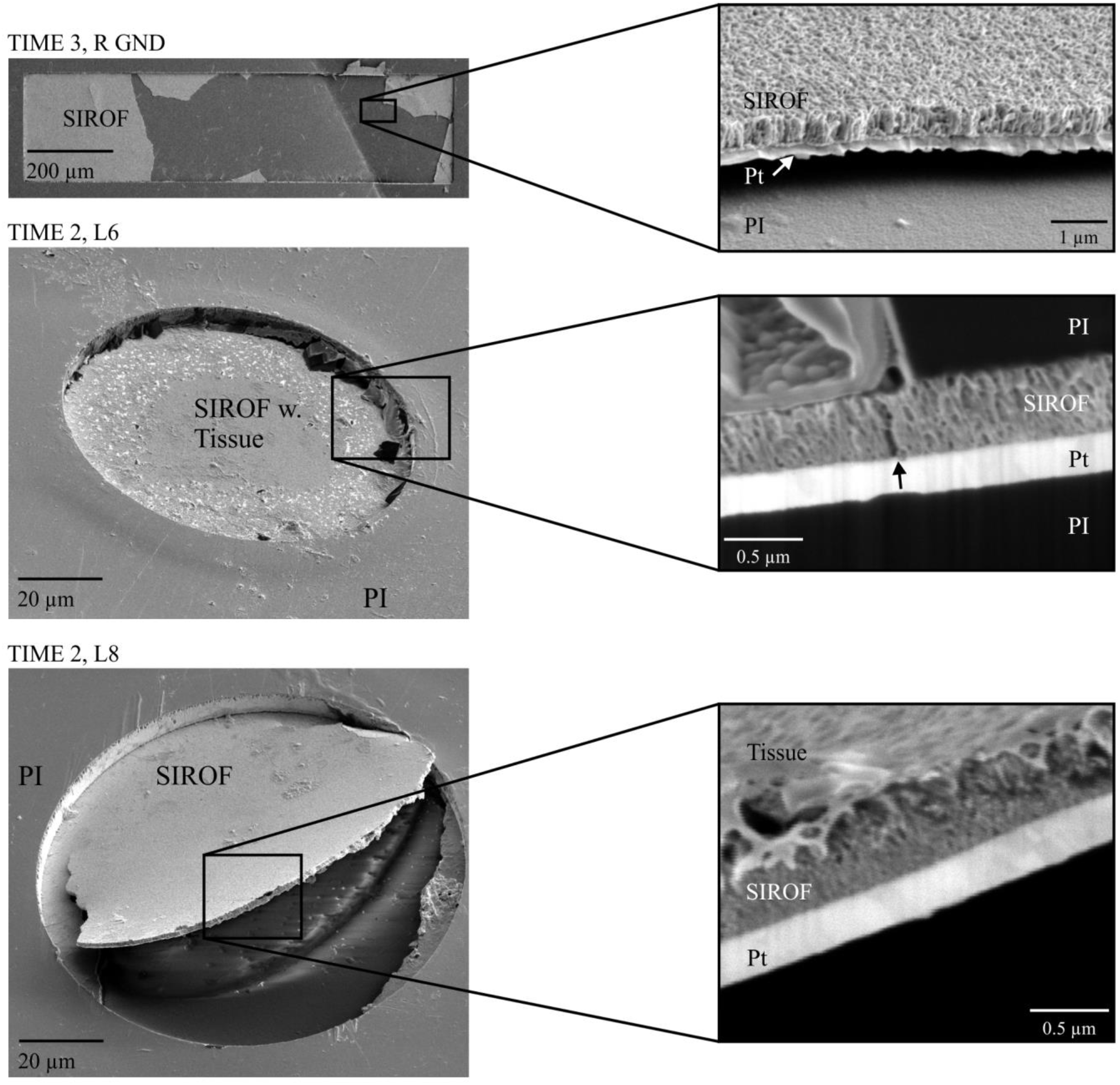
TIME-3H thin-film integrity after explantation. The right ground contact of the TIME 3 delaminated mostly. The detailed image depicts how the metallization is lifted. The stimulation contact site L6 of the TIME 2 stayed mechanically and electrically functional. Salt crystals dried at the PI-edge. Within the detailed image a slight crack formation in the SIROF metallization could be observed (arrow). The last contact on both sides (L8 and R8) was not connected electrically on none of the implants. Nevertheless, on the implant TIME 2 contact L8 the metallization delaminated completely.

### 3.3. Optical Analysis of Explanted RMIs (Rodent Trial)

Comparing the stimulation contact sites, 87.5 % of the TIME-3H sites were damaged in contrast to 0 % of the TIME-3H-SiC (Table 1). Three contacts of the TIME-3H used for extensive stimulation were analyzed and two failed. In the TIME-3H-SiC none of the 5 contacts used for stimulation failed. Concerning the ground contacts, all investigated grounds without SiC as adhesion promoter showed damage, even absence of the entire metallization (Figure 4), whereas the ground contacts with SiC underneath the platinum all were intact (Figure 4).

**Table 1.** Optically analyzed contact sites of explanted RMIs.

| El. type | Cont. type | Total av. | damaged |  | Used for stim | damaged |  |
| --- | --- | --- | --- | --- | --- | --- | --- |
|  |  |  | # | % |  | # | % |
| TIME-3H | Contact | 24 | 21 | 87.5 | 3 | 2 | 66.6 |
| TIME-3H-SiC | Contact | 24 | 0* | 0 | 5 | 0* | 0 |
| TIME-3H | GND | 4 | 4 | 100 | 4 | 4 | 100 |
| TIME-3H-SiC | GND | 3 | 0* | 0 | 3 | 0* | 0 |

\*For both, regular stimulation contact and ground contact of the TIME-3H-SiC one contact exhibited crack formation but no delamination.

**Figure 4.**
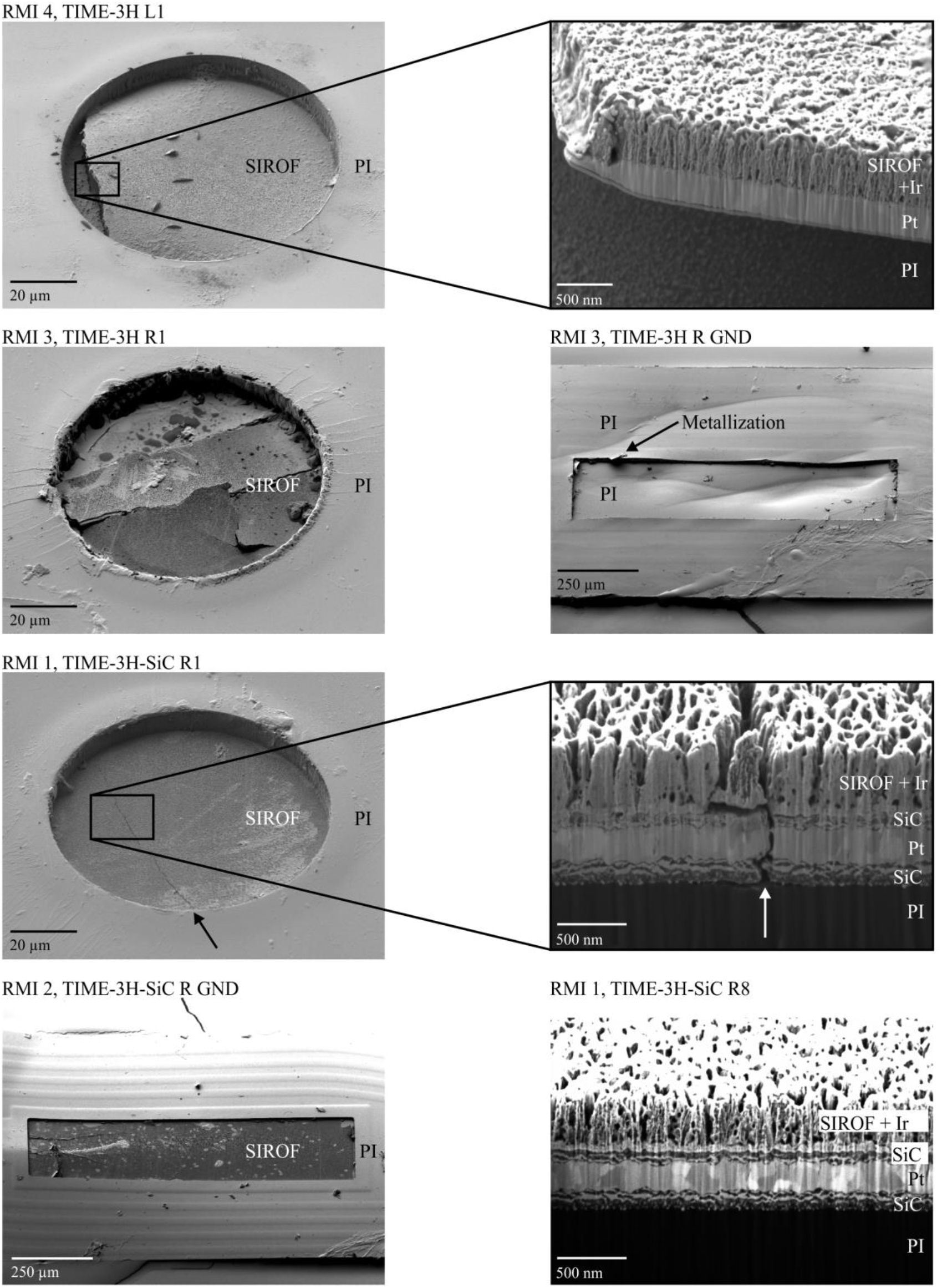
Thin-film electrode integrity after rodent animal trial. The upper four figures depict TIME-3H electrodes: RMI 4 exhibited delamination of the thin-film metallization compound on contact L1, which was used for stimulation. The same applied for contact R3 of RMI 3, including severe crack formation. The right ground of RMI 3 exhibited an entire delamination of the thin-film metallization. The lower four figures depict TIME-3H-SiC electrodes: On RMI 1 R1, which was used for stimulation, crack formation could be observed, but without delamination. The right ground contact of RMI 2 stayed entirely unimpaired. A cross-section of contact site R8 of the implant RMI 1 was acquired and unveiled the layer setup of the thin-film metallization, including a periodic pattern formation at the SiC-Pt-interface. Without adhesion promoting layer, 87.5 % of the stimulation contacts and 100 % of the ground contacts were damaged including crack formation and delamination. In contacts with SiC as adhesion layer, no delamination occurred. This was a significant improvement in terms of mechanical integrity.

TIME-3H thin-film electrodes delaminated in the RMIs, showing adhesion failure of the platinum-SIROF-sandwich and the PI, as well as crack formation with resulting delamination of the metallization (Figure 4, RMI 3 and RMI 4). In contrast, the contact sites of the TIME-3H-SiC stayed mechanically intact, despite slight crack formation, which did not lead to delamination (Figure 4, RMI 1, TIME-3H-SiC R1 arrow).

PI-based thin-film electrodes were analyzed from the bottom using optical microscopy with polarization filters. SiC exhibited a blue appearance (Figure 5, a and b). Cracks were visible as whitish lines (Figure 5 b, arrow). At the edge of the metallization, SiC exhibited a frizzy contour. As no interferences occurred, both contacts a and b in Figure 5 did not delaminate. On the contrary, the contacts without SiC (shiny appearance) showed interferences indicative of delamination due to height changes. Crack formation could be observed along the overlapping PI edge together with delamination (Figure 5 c, arrow). Also crack formation through the whole contact site causing delamination occurred (Figure 5 d, arrow).

**Figure 5.**
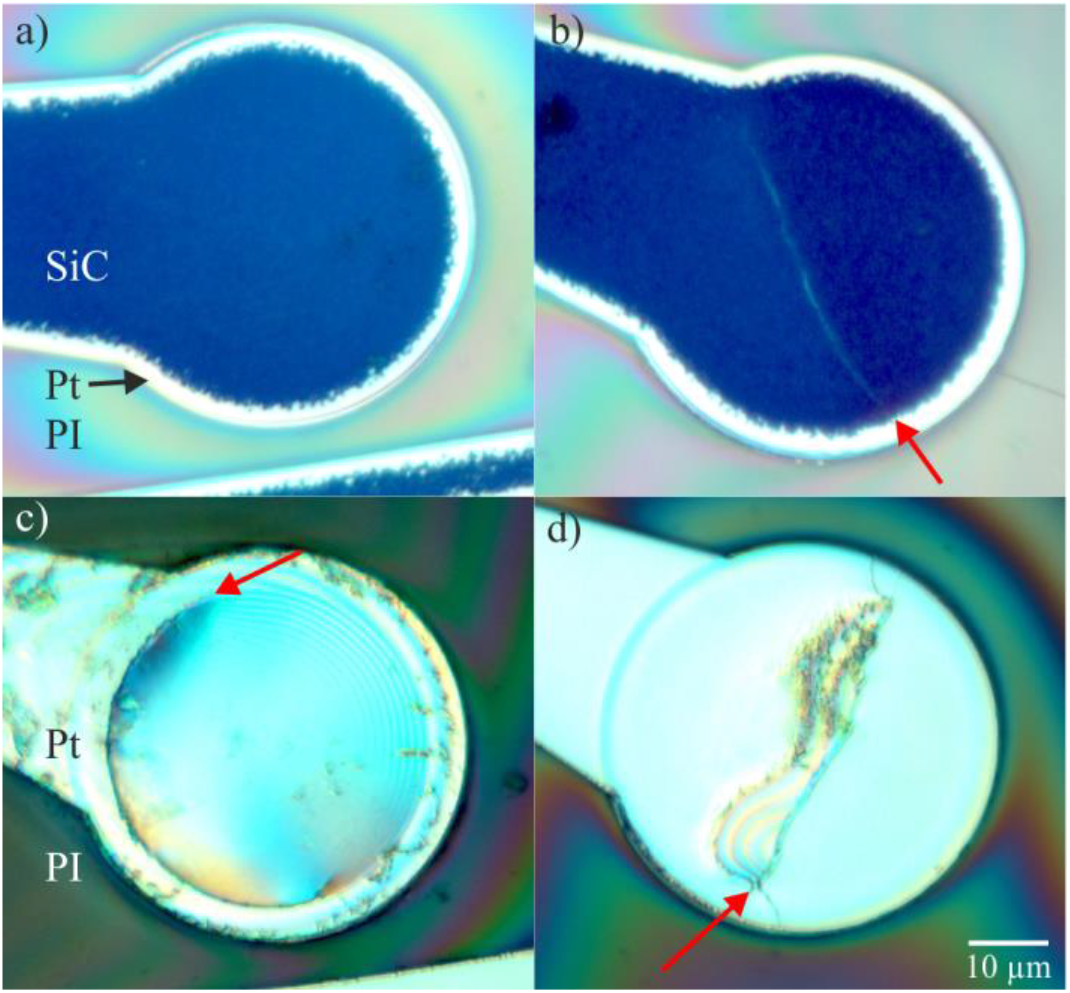
Microscopic view on the electrode backside through the bottom layer polyimide. A polarization filter was used to make delamination visible. SiC of the TIME-3H-SiC thin-film displayed a blue appearance (a and b). Crack formation could be observed as a whitish line through the metallization (arrow in b). The TIME-3H without SiC as adhesion promoting layer displayed a shiny appearance (c and d). Delamination was made visible with polarization filters, displaying interferences (c). Crack formation alongside the PI-overlap could be observed (arrow in c). Moreover, crack formation through the contact with partial delamination occurred (arrow in d).

## 4. Discussion

We have evaluated the performance of TIME thin-film electrodes designed and developed for a first-in-human clinical trial with a duration of 30 days. These electrodes proved good functionality for electrical stimulation in about 90 % of the stimulation contact sites. Since the ultimate goal is to apply such microimplants for years, careful analysis of changes in the electrode metal and the metal-polymer compound were done in a post-explantation study by means of light and scanning electron microscopy in addition to impedance spectroscopy measurements. The observations indicated that the adhesion between metals and polymer substrate and insulation needed to be improved for ensuring longer-term functionality.

The study was designed with two objectives for evaluating the used technology with respect to long-term stimulation applications and the technology transfer towards chronic clinical trials. We aimed to ascertain (1) how stable is the electrochemical transfer function of the stimulation sites over the implantation period and do stimulation thresholds stay within the chemical safe charge injection limit and (2) which lessons can be learned from explanted devices regarding the material interfaces as well as the surgical handling during explantation.

### 4.1. Electrical Characterization of the TIME Implants *in vivo* during the Human Clinical Trial

Electrode impedance characterization *in vivo* with one stimulation site versus one ground is a viable measure to evaluate the electrical functionality over the implantation time (Williams et al. 2007; Wurth et al. 2017) and to estimate the access resistance that originates from the tissue between the electrodes under test. It is a rough indirect estimate of the influence of foreign body reaction and fibrotic tissue encapsulation. In the human trial, 50 out of 56 stimulation contacts (89.3%) and 7 out of 8 ground contact sites (87.5%) were electrically functional after 30 days. In average, electrode impedance increased over the implantation period. Similar impedance increase over time was also found in other studies (Rossini et al. 2010) and was correlated with the developing fibrosis around the implant (Wurth et al. 2017). Stimulation threshold increased but remained well below the maximum safe charge injection limit (120 nC) of the iridium oxide coating of the stimulation contacts sites that is higher than for platinum, which limited the use of electrodes in a previous study (Rossini et al. 2010). Since the perceptions of the subject remained stable over time, we believe that the foreign body reaction around the TIMEs fixated the electrodes well inside the nerve in addition to the surgical fixation on the epineurium and prevented movements during the implantation time.

### 4.2. Optical Analysis of the TIME Electrodes from the Human Clinical Trial Post-Explantation

The explantation procedure at the end of the human study was performed prioritizing to minimize the time of the intervention and the potential damage to the subject nerve. Cables as well as the ceramic interconnector were tightly encapsulated due to the foreign body reaction, so a laborious dissection procedure was necessary to free cables and interconnect from this tissue to get explanted. Surgical needs and the protection of the patient during this intervention was more in focus than the integrity of the TIME devices. Therefore, several cuts were made to explant the implant in pieces and *ex vivo* evaluation of the TIME could not be made on complete devices but on fragments only. Especially the thin-film part of the TIME with the stimulation and the ground contact sites experienced mechanical forces during the explantation, that might have contributed to the mechanical delamination of the contact sites that had been already initiated during the implantation time due to loss of adhesion but did not deteriorate enough the electrical functionality.

Adhesion seemed to be at limit with the 30 days implantation period in the human trial with the chosen layer setup that had been validated in previous chronic animal experiments (Badia et al. 2011b; Harreby et al. 2015) before being transferred to the clinical setting. The milieu of the foreign body reaction with reactive oxygen species, pH value shifts and other factors released might contribute to material changes by incorporation of oxygen and hydrogen species into the platinum thin-film layer and the iridium oxide coating. This could lead to increased stress that was above the value of the adhesion force of the platinum-polyimide sandwich.

Another possibility for mechanical failure could be caused by contacts of filopodia of macrophages in the fibrotic tissue due to the foreign body response (la Oliva et al. 2018) and the rough iridium oxide surface of the electrode structure, that might be stronger than the adhesion between the platinum and the underlying PI. Cells react on nanostructures on the surface (Anselme et al. 2010) since they offer discrete attachment points. The topology of the nanostructures has a prevalent effect (Variola et al. 2009) and adhesion can be either increased (structures below 13 nm) or reduced (structures about 95 nm) (Anselme et al. 2010). Most investigations have been performed *in vitro* (Variola et al. 2009), with polymer surfaces (Patrito et al. 2007) or materials close to bone implants like titanium and aluminum oxide. Oxygen rich surfaces as well as large roughness tend to lead to larger fibroblast attachment (Patrito et al. 2007), but depending on the material, the larger initial cell attachment (Anselme et al. 2010; Ni et al. 2014) can eventually decrease fibroblast proliferation rate (Elter et al. 2011) on gold (Chapman et al. 2015) and platinum (Pennisi et al. 2009). Detailed analysis would be needed to solve the question of tissue adhesion on the electrode contact sites and the surrounding PI with samples that include part of the surrounding nerve and the implant which goes beyond the possibilities of this study since the major goal during the explantation procedure was to avoid damage to the remaining nerves of the patient.

As conclusion, adhesion forces of the PI-platinum compound have shown not to be sufficient for chronic implantation times and the benefit of adhesion layers to overcome this shortcoming had to be verified. The iridium oxide adhered well to the underlying platinum and did not cause any failure.

### 4.3. Outcome of Preclinical Study of TIME with Improved Metal-Polymer Adhesion

Integration of silicon carbide (SiC) was hypothesized by our group earlier as a potential solution to increase adhesion forces between platinum thin-film metallization and PI (Ordonez et al. 2012a; Ordonez et al. 2012b). TIME implants with a layer structure as used in the sub-chronic human implant study were compared to implants with additional SiC adhesion promotion layers underneath and above the platinum towards the PI in a rat model. We purposely designed a RMI for subcutaneous implant on the dorsum in order to allow that the two arms of TIME-3H and TIME-3H-SiC remained under the same conditions and could be used for the same extensive electrical stimulation. Despite the TIME is designed for implantation within the peripheral nerve, we preferred this subcutaneous implant for avoiding other tethering forces and leads breaks that are frequent in electrode implants in the rat limbs (la Oliva et al. 2018; Wurth et al. 2017). Nevertheless, the functional assessment of TIME chronic intraneural implant has been recently reported elsewhere (la Oliva et al. 2018).

Daily electrical stimulation for an accumulated average of 78 hours over around 1 month implant time corresponded to the number of stimulation pulses estimated for the intermittent stimulation within the human clinical trial of about two months (Raspopovic et al. 2014). Since both TIME versions without and with SiC as adhesion promoting layers were implanted side by side, they experienced identical environmental conditions in the same animal (four in total), thus ensuring comparability of results. While implants without SiC adhesion layers showed adhesion loss and delamination in 87.5% of stimulation sites and 100% of ground contacts, none (0%) of the platinum metallization of the stimulation sites and ground contacts with additional SiC coating showed signs of adhesion loss and delamination. While adhesion between platinum and PI alone is caused on physisorption with low binding energy and interlocking effects (Ordonez et al. 2012b; Ordonez 2013), SiC generates stronger covalent bonds to both, PI and platinum. The generation of platinum silicides mediates the chemisorption towards the platinum while direct bonds of the carbon atoms mediate adhesion to the carbon backbone of the PI.

## 5. Conclusion

Even though the electrical functionality of the stimulation sites and the clinical outcome of a sub-chronic first-in-human study with TIME implants was excellent (Oddo et al. 2016; Raspopovic et al. 2014), analysis of the explanted TIMEs revealed that the PI substrate and the platinum thin-film metallization underneath the iridium oxide coating did not strongly adhere to each other, no matter whether the electrode sites experienced electrical stimulation or not. Adhesion needs to be improved for potentially higher security and longer implantation periods. Silicon carbide (SiC) as adhesion promoter allowed for covalent bonds between SiC and platinum and PI, respectively, ensuring compound integrity and functionality even under intrinsic stresses during an *in vivo* study in a rat model with high stimulation regime. We believe that inclusion of such an adhesion layer can increase device stability of the thin-film part to the level that these devices can be transferred into chronic clinical trials.

## Supporting information

Supplementary Material for Manuscript Cvancara - Stability of Thin-Film Metallization in Flexible Stimulation Electrodes

## Author Contributions

Paul Čvančara designed and fabricated the RMIs and performed the analysis of the *in vivo* measurements and the optical analysis of the thin-film electrodes. He transferred the outcome into the manuscript and was writing the manuscript.

Tim Boretius designed and fabricated the TIME-3H implants and contributed to the manuscript with editing.

Thomas Stieglitz was the co-designer of TIME-3H and supervisor of this study, acted as scientific advisor and edited the manuscript.

Victor M López-Alvarez and Xavier Navarro designed the rodent model and performed the implants and stimulation protocol. Xavier Navarro also contributed to writing the manuscript.

Pawel Maciejasz, David Andreu, Jean-Louis Divoux and David Guiraud designed and manufactured the software and hardware of the stimulator and edited the manuscript. Pawel Maciejasz and David Guiraud acquired and preprocessed impedance data.

Stanisa Raspopovic, Francesco Petrini, Silvestro Micera, Giuseppe Granata, Eduardo Fernandez and Paolo M. Rossini performed the human clinical trial and edited the manuscript.

Winnie Jensen gave scientific advice in the clinical study and contributed in manuscript proof-reading.

Ken Yoshida gave scientific advice in the clinical study and contributed in manuscript proof-reading.

The manuscript was written through contributions of all authors. All authors have given approval to the final version of the manuscript.

## Acknowledgement

This work was supported by grants TIME (CP-FP-INFSO 224012) and EPIONE (FP7-HEALTH-2013-INNOVATIO-1 602547) from the European Commission (EC). The authors would like to thank the whole working group – especially those who are not on the list of authors – of the TIME and the EPIONE projects for their contributions and collaboration in the projects. Moreover, the authors would like to thank the KNMF at Karlsruhe Institute of Technology (especially Dr. Sabine Schlabach) for the granted working hours at the FIB (ID 2014-012004429).

## References

Anselme, K., Davidson, P., Popa, A.M., Giazzon, M., Liley, M., Ploux, L., 2010. Acta Biomater 6 (10), 3824–3846.

Badia, J., Boretius, T., Andreu, D., Azevedo-Coste, C., Stieglitz, T., Navarro, X., 2011a. J Neural Eng 8 (3), 36023.

Badia, J., Boretius, T., Pascual-Font, A., Udina, E., Stieglitz, T., Navarro, X., 2011b. IEEE Trans Biomed Eng 58 (8), 2324–2332.

Boretius, T., 2013. TIME. A Transverse Intrafascicular Multichannel Electrode. Zugl.: Freiburg im Breisgau, Univ., Diss., 2013. Der Andere Verlag, Uelvesbüll.

Boretius, T., Badia, J., Pascual-Font, A., Schuettler, M., Navarro, X., Yoshida, K., Stieglitz, T., 2010. Biosens Bioelectron 26 (1), 62–69.

Boretius, T., Stieglitz, T., 2012, 279–282.

Boretius, T., Yoshida, K., Badia, J., Harreby, K., Kundu, A., Navarro, X., Jensen, W., Stieglitz, T., 2012. A transverse intrafascicular multichannel electrode (TIME) to treat phantom limb pain — Towards human clinical trials, in: 2012 4th IEEE RAS & EMBS International Conference on Biomedical Robotics and Biomechatronics (BioRob). IEEE, pp. 282–287.

Branner, A., Stein, R.B., Fernandez, E., Aoyagi, Y., Normann, R.A., 2004. IEEE Trans Biomed Eng 51 (1), 146–157.

Campbell, P.K., Jones, K.E., Huber, R.J., Horch, K.W., Normann, R.A., 1991. IEEE Trans Biomed Eng 38 (8), 758–768.

Chapman, C.A.R., Chen, H., Stamou, M., Biener, J., Biener, M.M., Lein, P.J., Seker, E., 2015. ACS Appl Mater Interfaces 7 (13), 7093–7100.

Cogan, S.F., 2008. Annu Rev Biomed Eng 10, 275–309.

Elter, P., Weihe, T., Lange, R., Gimsa, J., Beck, U., 2011. European biophysics journal : EBJ 40 (3), 317–327.

Fiedler, E., Ordonez, J.S., Stieglitz, T., 2013. Biomed Tech (Berl).

Harreby, K.R., Kundu, A., Yoshida, K., Boretius, T., Stieglitz, T., Jensen, W., 2015. Artificial organs 39 (2), E36-48.

Hassler, C., Boretius, T., Stieglitz, T., 2011. J Polym Sci B Polym Phys 49 (1), 18–33.

Kundu, A., Harreby, K.R., Yoshida, K., Boretius, T., Stieglitz, T., Jensen, W., 2014. IEEE Trans Neural Syst Rehabil Eng 22 (2), 400–410.

la Oliva, N. de, Navarro, X., Del Valle, J., 2018. Journal of biomedical materials research. Part A 106 (3), 746–757.

Naples, G.G., Mortimer, J.T., Scheiner, A., Sweeney, J.D., 1988. IEEE Trans Biomed Eng 35 (11), 905–916.

Navarro, X., Krueger, T.B., Lago, N., Micera, S., Stieglitz, T., Dario, P., 2005. J. Peripher. Nerv. Syst. 10 (3), 229–258.

Ni, S., Sun, L., Ercan, B., Liu, L., Ziemer, K., Webster, T.J., 2014. Journal of biomedical materials research. Part B, Applied biomaterials 102 (6), 1297–1303.

Oddo, C.M., Raspopovic, S., Artoni, F., Mazzoni, A., Spigler, G., Petrini, F.M., Giambattistelli, F., Vecchio, F., Miraglia, F., Zollo, L., Di Pino, G., Camboni, D., Carrozza, M.C., Guglielmelli, E., Rossini, P.M., Faraguna, U., Micera, S., 2016. Elife 5, e09148.

Ordonez, J.S., 2013. Miniaturization of Neuroprosthetic Devices and the Fabrication of a 232-Channel Vision Prosthesis with a Hermetic Package. Der Andere Verlag, Uelvesbüll.

Ordonez, J.S., Boehler, C., Schuettler, M., Stieglitz, T., 2012a. Conf Proc IEEE Eng Med Biol Soc 2012, 5134–5137.

Ordonez, J.S., Schuettler, M., Boehler, C., Boretius, T., Stieglitz, T., 2012b. MRS Bull 37 (06), 590–598.

Patrito, N., McCague, C., Norton, P.R., Petersen, N.O., 2007. Langmuir : the ACS journal of surfaces and colloids 23 (2), 715–719.

Pennisi, C.P., Sevcencu, C., Dolatshahi-Pirouz, A., Foss, M., Hansen, J.L., Larsen, A.N., Zachar, V., Besenbacher, F., Yoshida, K., 2009. Nanotechnology 20 (38), 385103.

Raspopovic, S., Capogrosso, M., Petrini, F.M., Bonizzato, M., Rigosa, J., Di Pino, G., Carpaneto, J., Controzzi, M., Boretius, T., Fernandez, E., Granata, G., Oddo, C.M., Citi, L., Ciancio, A.L., Cipriani, C., Carrozza, M.C., Jensen, W., Guglielmelli, E., Stieglitz, T., Rossini, P.M., Micera, S., 2014. Sci Transl Med 6 (222), 222ra19.

Rossini, P.M., Micera, S., Benvenuto, A., Carpaneto, J., Cavallo, G., Citi, L., Cipriani, C., Denaro, L., Denaro, V., Di Pino, G., Ferreri, F., Guglielmelli, E., Hoffmann, K.-P., Raspopovic, S., Rigosa, J., Rossini, L., Tombini, M., Dario, P., 2010. Clin Neurophysiol 121 (5), 777–783.

Rousche, P.J., Normann, R.A., 1998. J. Neurosci. Methods 82 (1), 1–15.

Rubehn, B., Bosman, C., Oostenveld, R., Fries, P., Stieglitz, T., 2009. J Neural Eng 6 (3), 36003.

Rubehn, B., Stieglitz, T., 2010. Biomaterials 31 (13), 3449–3458.

Stieglitz, T., Beutel, H., Meyer, J.-U., 2000a. J Intell Mater Syst Struct 11 (6), 417–425.

Stieglitz, T., Beutel, H., Schuettler, M., Meyer, J.-U., 2000b. Biomed Microdevices 2 (4), 283–294.

Variola, F., Vetrone, F., Richert, L., Jedrzejowski, P., Yi, J.-H., Zalzal, S., Clair, S., Sarkissian, A., Perepichka, D.F., Wuest, J.D., Rosei, F., Nanci, A., 2009. Small (Weinheim an der Bergstrasse, Germany) 5 (9), 996–1006.

Williams, J.C., Hippensteel, J.A., Dilgen, J., Shain, W., Kipke, D.R., 2007. J Neural Eng 4 (4), 410–423.

Wurth, S., Capogrosso, M., Raspopovic, S., Gandar, J., Federici, G., Kinany, N., Cutrone, A., Piersigilli, A., Pavlova, N., Guiet, R., Taverni, G., Rigosa, J., Shkorbatova, P., Navarro, X., Barraud, Q., Courtine, G., Micera, S., 2017. Biomaterials 122, 114–129.

