## Supplementary Material for Manuscript Cvancara - Stability of Thin-Film Metallization in Flexible Stimulation Electrodes for "Stability of Thin-Film Metallization in Flexible Stimulation Electrodes: Analysis and Improvement of *in vivo* Performance"

### 1. Materials and Methods

#### 1.1. Design of the TIME thin-film layout

The thin-film electrode arrays of the TIME implants were fabricated using standard processing of microelectromechanical systems (MEMS). It is in the nature of standard MEMS processing that the final product is a 2D-structure. Considering the implantation procedure of the TIME (Krähenbühl et al. 2017), it might happen, that the electrode sites remain between two fascicles (Kundu et al. 2014). Given this anatomical situation, it is of advantage having contact sites on both sides of the implant, in order to stimulate the fascicles left and right to it. A simple way to realize electrical contacts on both sides of a 2D-substrate, is to fold it. Double-side arrangement requires technologies that are much more sophisticated and often result in much lower yield (Stieglitz 2001). In case of the TIME implant, a U-shaped design is used (Figure S1 a), which is folded in the middle and results in a L-shape (Boretius et al. 2012). This L-shape can be divided into three parts. First, the transition part which contains a special structure to interconnect electrically and mechanically the polyimide thin-film with a following assembly setup containing a screen-printed ceramic. Microflex interconnection technique (MFI) (Stieglitz et al. 2000) is used as a robust and reliable technique to assemble a thin-film ribbon with a long helically wound cable (Figure S1 a). The MFI structure design contains a 60 µm hole for the mechanical connection, surrounded by a platinum ring of 20 µm width for the electrical connection. Second, the ribbon part contains all tracks which lead from the transition part to the stimulation contact sites. Moreover, it contains the large area ground contact sites. Third, the electrode part consists of the active strip with the stimulation contact sites incorporated and after folding a loop is shaped out of the active strip. A loop of surgical thread is incorporated within the PI loop together with an integrated surgical needle to pull the implant transversally through the nerve during implantation. Flaps with holes are integrated in the design to allow surgical fixation of the TIME to the epineurium of the nerve. The holes are surrounded by metal rings for better visual detection. The flaps are arranged on the ribbon part that goes in parallel to the nerve axis after implantation.

The detailed implantation procedure is described elsewhere (Boretius et al. 2012; Raspopovic et al. 2014). In general, the front part of the TIME design (active part) is pulled with the incorporated needle and suture during the implantation through the nerve. The active sites for stimulation have to lay inside the nerve. The fixation rings and the loop are sutured to the surrounding epineurium to prevent movements of the implant until it is fixed by a fibrotic sheath as result of the foreign body reaction.

The version TIME-3H, used in the first-in-human study sub-chronically, contained 18 channels, of which 16 belonged to the stimulation contact sites (only 14 connected, due to the limited amount of channels within the connector) and two ground contacts. After folding, the electrode displayed seven connected stimulation contact sites on the left and seven on the right, named L1 to L7 and R1 to R7 respectively. The same applied for the two ground contacts, L GND and R GND. The stimulation contact sites had a circular shape with a diameter of d = 80 µm. The pitch between the active sites was 0.4 mm, resulting in an active strip length of 2.48 mm (Figure S1, blue box). The large ground sites exhibited a rectangular shape with an area of 1.00 x 0.25 mm² each (Figure S1, red box). Platinum tracks and pads were sandwiched between polyimide as substrate and insulation layer. Platinum tracks were completely surrounded by the polyimide (details in (Boretius et al. 2012) and below). At the active sites and ground contacts, the platinum layer was coated by iridium, subsequently covered by a sputtered iridium oxide film (SIROF) (Figure S1 b, left) and opened via reactive ion etching (RIE).

The TIME-3H-SiC used the same design, but with a different layer setup in order to investigate SiC as adhesion promoting layer (Figure S1 b, right).

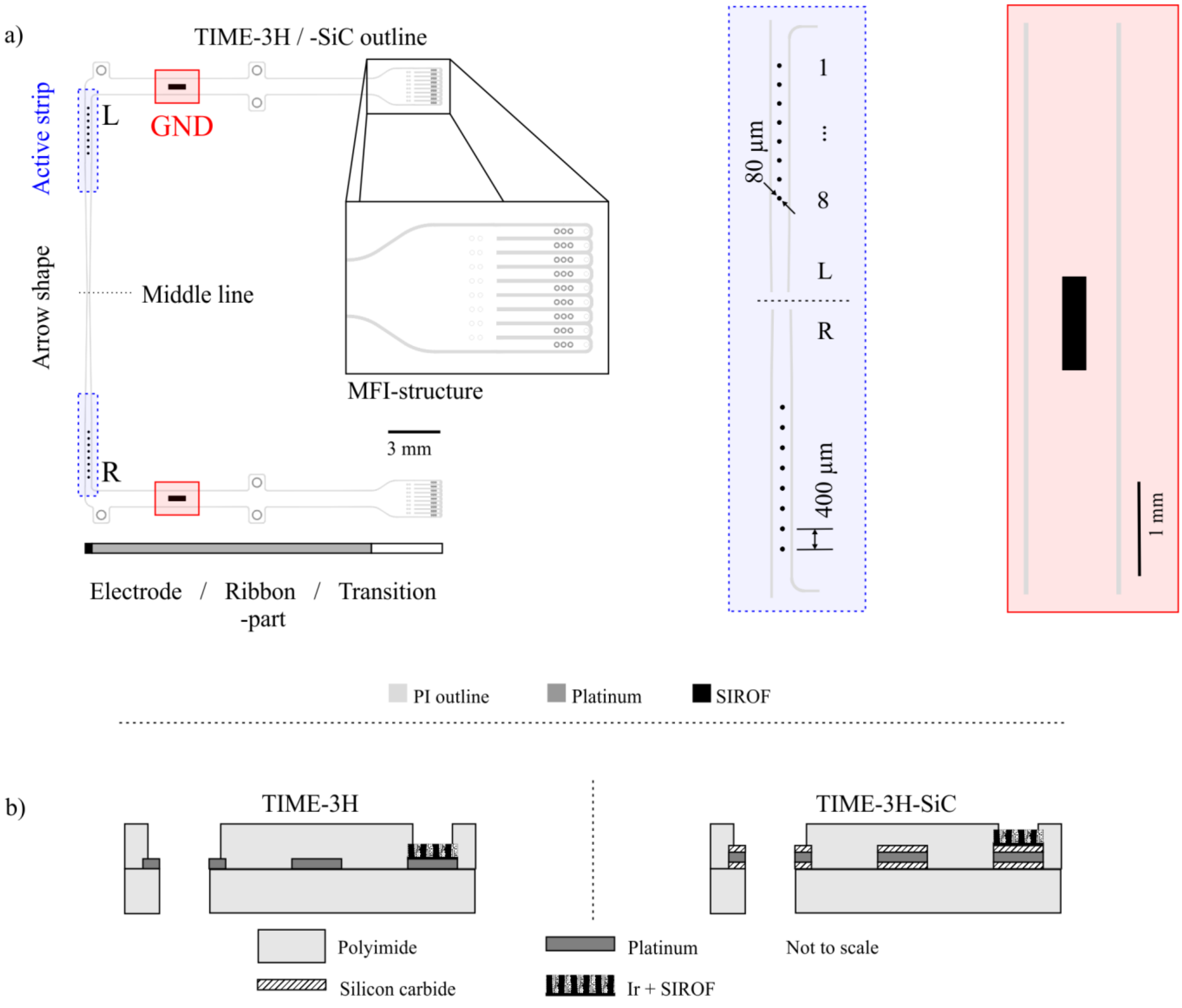


**Figure S1.** Overview of the TIME-3H design. The thin-film electrode consists of two mirrored sides with 8 stimulation contacts and one ground each (a). The TIME-3H-SiC used the same design but a different layer setup (b).

#### 1.2. Cleanroom fabrication of thin-film electrodes

The micro-fabrication of the various thin-film electrodes was conducted in clean room environment class 1000 with standard photolithography and MEMS processes (Figure S2).

Both thin-film electrode versions used polyimide (PI, U-Varnish S, UBE Industries, LTD., Tokyo, Japan) as substrate and insulation material. A 5 µm film of PI was spin-coated onto a

4"-silicon wafer (Figure S2 a) and imidized at 450 °C in a nitrogen atmosphere in a furnace (YES-459PB6-2PE-CP, Yield Engineering Systems Inc., San Jose, CA, USA).

In case of the TIME-3H 1.4 µm of a high-resolution image reversal resist (AZ 5214E, MicroChemicals GmbH, Ulm, Germany) was applied via spin coating. The resist served as deposition mask for the succeeding track metallization. After exposure and development of the resist, an $O_2$-plasma activation step in a reactive ion etching chamber (RIE, RIE Multiplex, STS Surface Technology Systems plc, Newport, UK) to increase reactivity of the PI surface was conducted. Subsequently, a 300 nm layer of platinum was sputter deposited (Pt, Leybold Univex 500, Leybold Vacuum GmbH, Cologne, Germany) (Figure S2 b). To remove excessive material and the resist, lift-off with acetone and isopropyl alcohol was performed. Afterwards, a further image reversal resist defining the stimulation contact sites and ground contacts was spin coated, exposed and developed. 100 nm of iridium and subsequently 800 nm iridium oxide (SIROF) were sputter deposited (Ir, Leybold Univex 500, Leybold Vacuum GmbH, Cologne, Germany) (Figure S2 c), to ensure high charge injection capacities (Cogan et al.; Weiland and Anderson 2000). Following another $O_2$-plasma surface activation, a second layer of PI with a thickness of approximately 5 µm was spin coated to insulate the metallization (Figure S2 d). A positive resist (AZ 9260, MicroChemicals GmbH, Ulm, Germany) was utilized as etching mask. The perimeters and openings were realized using RIE in an oxygen plasma.

In case of the TIME-3H-SiC – in order to evaluate SiC as adhesion promoter – a 50 nm layer of SiC was deposited using plasma-enhanced chemical vapor deposition (PECVD, PC310 reactor, STS Surface Technology Systems plc, Newport, UK) before evaporation of a 300 nm layer of platinum (Leybold Univex 500, Leybold Vacuum GmbH, Cologne, Germany) (Figure S2 b). Directly afterwards, another adhesion promoting layer of 50 nm SiC was deposited via

PECVD on the platinum (Figure S2 b1). All further fabrication process steps were the same for both versions of TIME-3H.

Both, the TIME-3H thin-films used for the human clinical trials and the TIME-3H-SiC for adhesion promotion validation were investigated with special cytotoxicity samples (Stieglitz et al. 2011) (run in parallel with the same process) according to the ISO 10993 for cytotoxicity testing. Direct contact and extract tests were performed compliant with the ISO 10993-5 with L929 mouse fibroblasts. Additionally, further direct contact and extract tests were performed with the human nerve cell line Kelly and the human muscle cell line A673. All samples passed the tests and no objections were claimed.

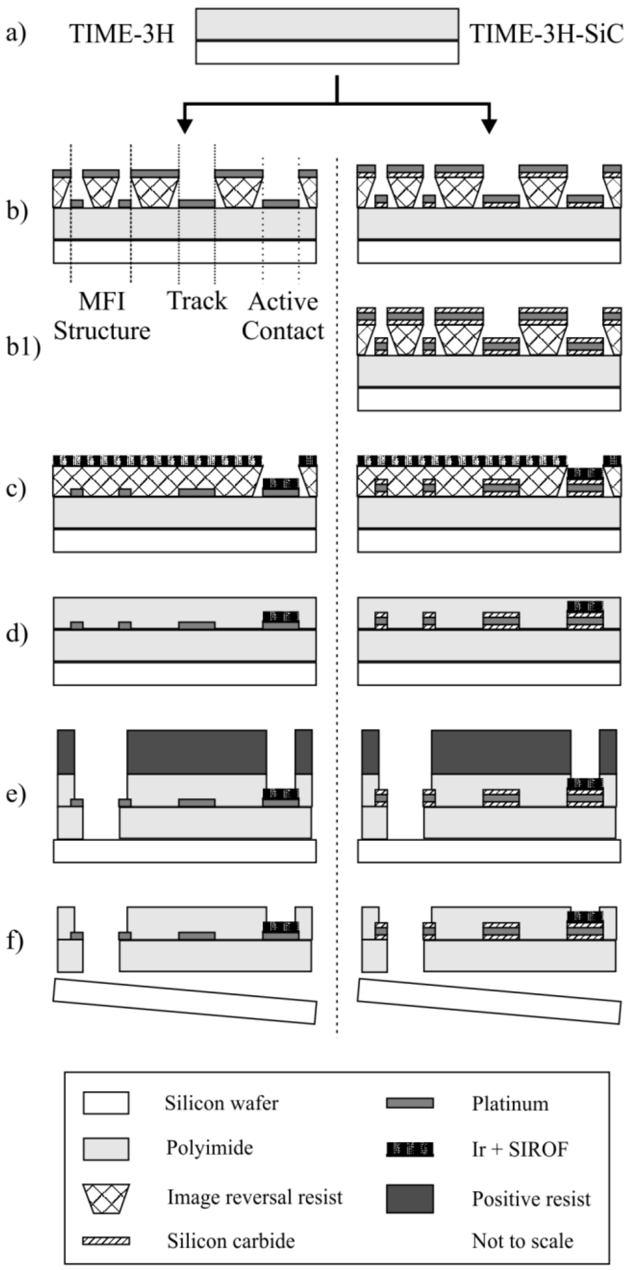


**Figure S2.** Fabrication process of TIME thin-film electrodes. Process variations for the TIME-3H (left) and TIME-3H-SiC are depicted. After spin-coating and imidizing a 5 µm layer of PI (a), an image reversal resist was applied as structuring mask. Next, 300 nm of platinum were sputter deposited (b). In case of the TIME-3H-SiC 50 nm SiC were applied via PECVD and 300 nm platinum were evaporated. Afterwards, 40 nm SiC were created (b1). Following another resist, iridium and SIROF were sputter deposited on the active and ground contact sites (c). For electrical insulation a second layer of 5 µm PI was spin-coated and imidized (d), before opening the perimeters and the contact sites with RIE (e).

##### 1.3. Electrical characterization of implants *in vitro*

In order to characterize and compare the electrochemical properties of the TIMEs and the RMI – with both electrode types – the devices underwent an analysis with electrochemical impedance spectroscopy (EIS) *in vitro* before implantation. The characterization was performed with a frequency analyzer and a potentiostat (SI 1260 & SI 1287, Solartron Analytical, Farnborough, UK) in phosphate-buffered saline (PBS) solution with a frequency range of 100 kHz to 1 Hz and a voltage amplitude of 10 mV. The three-electrode setup consisted of an Ag/AgCl reference (3M), a large area platinum counter and the respective active and ground contact sites of the TIME as working electrode.

##### 1.4. Electrical characterization of the TIMEs *in vivo* (human)

The applied current for impedance estimation did not evoke any sensation in the patients' perception. Five stimuli were used for each contact assessment. The first one was discarded and the value of the estimated impedance was averaged on the four remaining pulses. The measurement was discarded if the current was not delivered properly and thus the contact was considered as open circuit without estimation of the impedance.

Measurements were performed for each stimulation contact site versus both, L GND and R GND separately. The ground contacts were characterized vice versa against each other. 128 measurements (4 TIMEs x 4 sides versus ground x 8 contacts) were performed within one session in less than 15 minutes, leading to 512 pulses to process.

#### 2. Results

##### 2.1. Electrical characterization of implants *in vitro*

Electrochemical characterization of the different implants and the different layer setups prior to implantation was performed using EIS. Electrode sites showed the expected high pass behavior (Figure S3), with differences in TIME-3H and RMI, but not much variability within the batches of the devices (Table S1).

The contacts of the TIME-3H implant exhibited an impedance of about 3 kΩ higher than the contacts of the RMI at a frequency of f = 1 kHz. On the different thin-film electrode versions of the RMI on the other hand, the impedances differed only with approximately 0.6 kΩ. The same was true for the ground contact sites. The impedance of the TIME-3H implant was about 0.3 kΩ higher than the equal ground impedances of both thin-film types of the RMI. The phase shift and cut-off frequencies exhibited the same behavior. Both, active contact sites and ground contacts of the TIME implants were higher than the almost equal properties of the different thin-film electrodes of the RMIs.

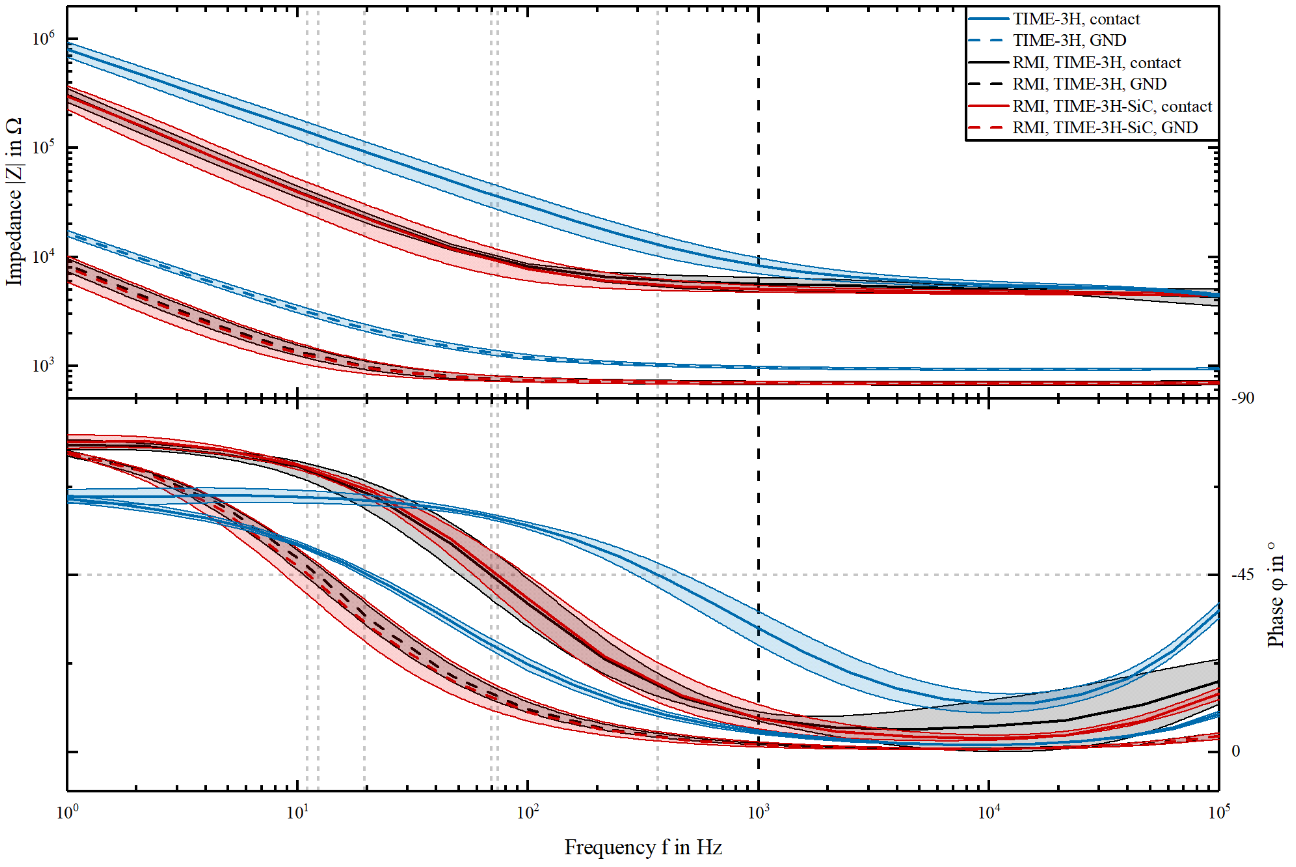


**Figure S3.** Electrochemical characterization of the TIMEs and RMIs (n = 5 per contact site). The TIME had a shifted cut-off-frequency, higher impedances and shifted phase angle compared to the RMIs, as the implants were not hydrated before characterization and had a higher access resistance due to the high resistivity of the helically wound cable. There was nearly no difference between the different thin-film electrode types on the RMIs.

**Table S1.** Electrochemical properties of the various implants *in vitro*.

| Contact type | Type (diameter) | Impedance \|Z\| in kΩ @ 1 kHz | Phase φ in ° @ 1 kHz | Cut-off frequency $f_{cut}$ in Hz |
|---|---|---|---|---|
| Stimu-lation | TIME-3H, (80 µm) | 8.3 ± 1.41 | -31.4 ± 4.4 | 362.0 |
| | TIME-3H, RMI, (80 µm) | 5.6 ± 0.81 | -8.5 ± 0.6 | 69.6 |
| | TIME-3H-SiC, RMI, (80 µm) | 5.0 ± 0.33 | -8.5 ± 3.1 | 74.0 |
| GND | TIME-3H | 1.0 ± 0.02 | -5.1 ± 0.5 | 20.0 |
| | TIME-3H, RMI | 0.7 ± 0.02 | -1.9 ± 0.2 | 12.3 |
| | TIME-3H-SiC, RMI | 0.7 ± 0.02 | -1.8 ± 0.6 | 11.0 |

The electrochemical properties of the electrode sites on devices that have been manufactured in two different clean-room runs for the human clinical trial and the rodent model implants (RMI) varied, showing higher impedance magnitude and cut-off frequencies in the human devices. Hydration directly before characterization results in the lower impedance magnitude and cut-off frequency of the RMI devices (Boretius 2013; Boretius and Stieglitz 2012). The use of silicon carbide as adhesion promoter did not influence the electrochemical parameters as shown in the RMI investigations. TIME-3H and TIME-3H-SiC exhibited nearly identical electrochemical properties in the RMI study.

#### 2.2. Electrical characterization of the TIMEs *in vivo* (human)

The impedances of the TIME 1 measured versus the R GND exhibited similar properties like the measurements versus L GND (Figure S4, upper left). TIME 2 versus R GND exhibited similar properties compared to TIME 2 versus L GND. The only difference was on day 17, with a 30 kΩ higher impedance including 40 kΩ higher standard deviation (Figure S4, upper right). In case of the implant TIME 3, on day 30 both, the contact sites impedances and the R GND contact impedance were not measurable (Figure S4, lower left). TIME 4 exhibited similar properties compared to TIME 4 versus L GND (Figure S4, lower right). In total, the mean impedance of all stimulation contact sites increased from 32 kΩ ± 49 kΩ to 85 kΩ ± 36 kΩ, measured versus the R GND.

Same statistical analysis was applied to the impedance data acquired using the R GND for measurement. The impedances behaved similar like above, using the L GND for acquisition (Figure S4).

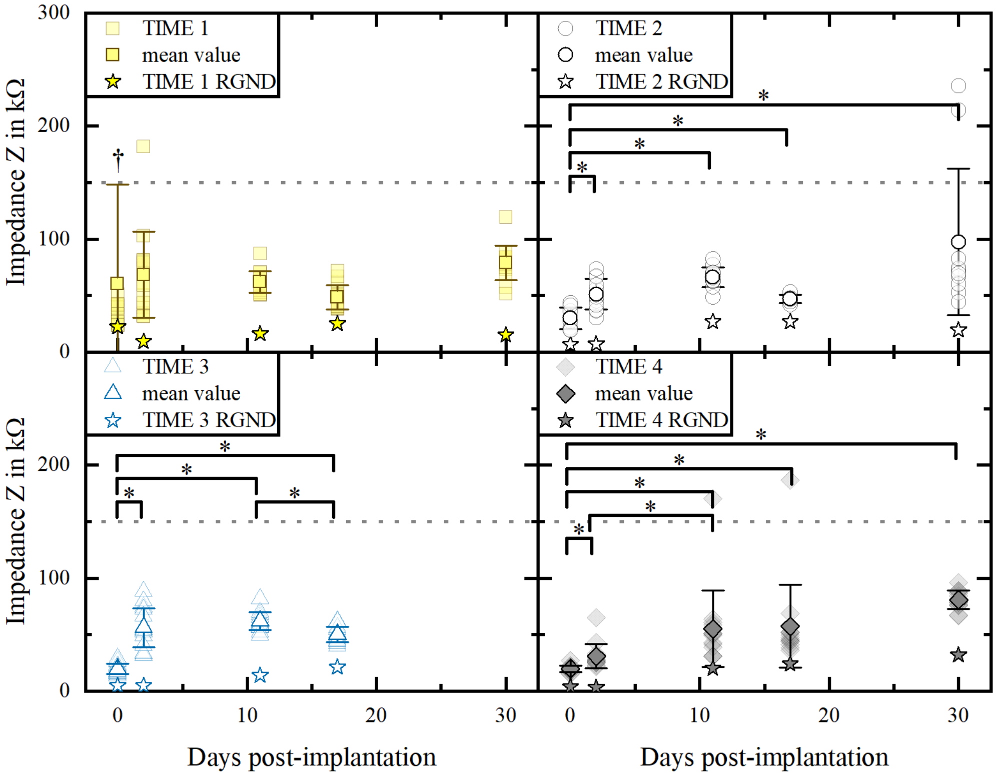


**Figure S4.** Impedance progression of four sub-chronically implanted TIME-3H implants. The impedances of the single active contact sites were measured versus the R GND of the respective implant. The measurements of each contact site are transparent (†: TIME 1, day 0, one impedance value at 365 kΩ), whereas the mean values are in bright color. The stars indicate the impedance of R GND, measured versus L GND. Electrical non-functionality is marked at 150 kΩ. The impedances of TIME 1 were statistically not different between each time point (linear mixed effect model; $p = 0.4834$). There was statistically significant difference between the impedances of specific days (especially day 0 compared to the other days) of the implants TIME 2, TIME 3 and TIME 4 (paired-sample Wilcoxon signed rank test with Bonferroni correction; $p < 0.01$).

#### 2.3. Optical analysis of explanted TIME implants (human)

Cross-sections of a newly fabricated, pristine PI-based TIME-3H thin-film electrode were acquired using FIB-SEM for comparison (Figure S5). The upper metallization consisted of the typical “cauliflower”-like topology of SIROF with a thickness of approximately 500 nm.

Underneath, a 100 nm thick layer of iridium was found on 300 nm platinum. The metallization conformly adhered on the PI substrate.

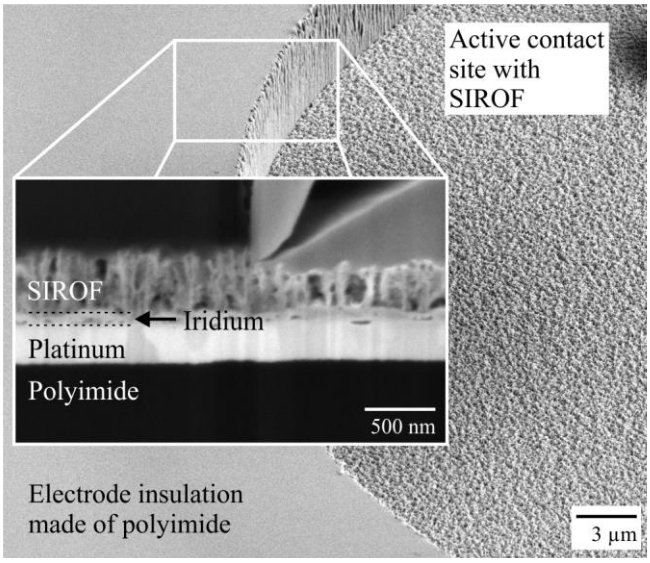


**Figure S5.** Pristine TIME-3H thin-film electrode. An active contact site with SIROF metallization was cut by FIB to gather a cross-sectional view with a SEM.

References

Boretius, T., 2013. TIME. A Transverse Intrafascicular Multichannel Electrode. Zugl.: Freiburg im Breisgau, Univ. , Diss., 2013. Der Andere Verlag, Uelvesbüll.

Boretius, T., Stieglitz, T., 2012, 279–282.

Boretius, T., Yoshida, K., Badia, J., Harreby, K., Kundu, A., Navarro, X., Jensen, W., Stieglitz, T., 2012. A transverse intrafascicular multichannel electrode (TIME) to treat phantom limb pain — Towards human clinical trials, in: 2012 4th IEEE RAS & EMBS International Conference on Biomedical Robotics and Biomechatronics (BioRob). IEEE, pp. 282–287.

Cogan, S.F., Plante, T.D., Ehrlich, J. Sputtered iridium oxide films (SIROFs) for low-impedance neural stimulation and recording electrodes, pp. 4153–4156.

Krähenbühl, S.M., Čvančara, P., Stieglitz, T., Bonvin, R., Michetti, M., Flahaut, M., Durand, S., Deghayli, L., Applegate, L.A., Raffoul, W., 2017. Medicine 96 (29), e7528.

Kundu, A., Harreby, K.R., Yoshida, K., Boretius, T., Stieglitz, T., Jensen, W., 2014. IEEE Trans Neural Syst Rehabil Eng 22 (2), 400–410.

Raspopovic, S., Capogrosso, M., Petrini, F.M., Bonizzato, M., Rigosa, J., Di Pino, G., Carpaneto, J., Controzzi, M., Boretius, T., Fernandez, E., Granata, G., Oddo, C.M., Citi, L., Ciancio, A.L., Cipriani, C., Carrozza, M.C., Jensen, W., Guglielmelli, E., Stieglitz, T., Rossini, P.M., Micera, S., 2014. Sci Transl Med 6 (222), 222ra19.

Stieglitz, T., 2001. Sens Actuators A Phys 90 (3), 203–211.

Stieglitz, T., Beutel, H., Meyer, J.-U., 2000. J Intell Mater Syst Struct 11 (6), 417–425.

Stieglitz, T., Schuettler, M., Rubehn, B., Boretius, T., Badia, J., Navarro, X., 2011, 529–533.

Weiland, J.D., Anderson, D.J., 2000. IEEE Trans Biomed Eng 47 (7), 911–918.
